# High pathogenicity avian influenza virus H5N1 (clade 2.3.4.4b) drives mass mortality in Eurasian crane (*Grus grus*) populations in Germany, 2025

**DOI:** 10.64898/2025.12.08.692485

**Authors:** Anne Günther, Christof Herrmann, Julia Sehl-Ewert, Simon Piro, Ann Kathrin Ahrens, Sten Calvelage, Anne Pohlmann, Martin Beer, Timm Harder

## Abstract

In autumn 2025, an unprecedented mass mortality event was observed among the western migrating subpopulation of Eurasian cranes (*Grus grus*) in Germany. Systemic infection with highly pathogenic avian influenza virus H5N1, clade 2.3.4.4b, genotype DI.2.1, was identified as the cause of acute death. The gregarious behaviour of cranes at feeding and resting sites likely has contributed to the rapid and massive dissemination of viruses within the crane population.

## Main text

Eurasian cranes (*Grus grus*) migrate along eastern, central and western European flyways. Mass mortalities caused by goose-Guangdong (gs/GD)-like high pathogenicity avian influenza viruses (HPAIV) of subtype H5 have shown this species’ susceptibility on the eastern and central flyway in the Near East, 2021/22 and 2024/25 [1], and in Eastern Europe, 2023/24 [2] and 2023/25 [3]. Cranes on the western flyway were spared from severe outbreak events so far, despite the ongoing HPAI enzootic in Europe. This changed in October 2025, when widespread deaths were detected in Germany, and thereafter, in France and Spain.

Each year, more than 420,000 cranes migrate through Germany, using approximately 300 roost sites. Scandinavian birds typically roost along the coast in Mecklenburg-Western Pomerania, while cranes from Finland, Poland, and the Baltic region prefer inland staging sites such as Lake Galenbeck, the Müritz region, Rhin-Havelluch or the Berga/Kelbra reservoir (Figure 1B, #1-4). Via the roosting region Diepholzer fen (Figure 1B, #5) in northwestern parts of Germany, they continue to wintering areas in France and Spain.

**Figure 1.**
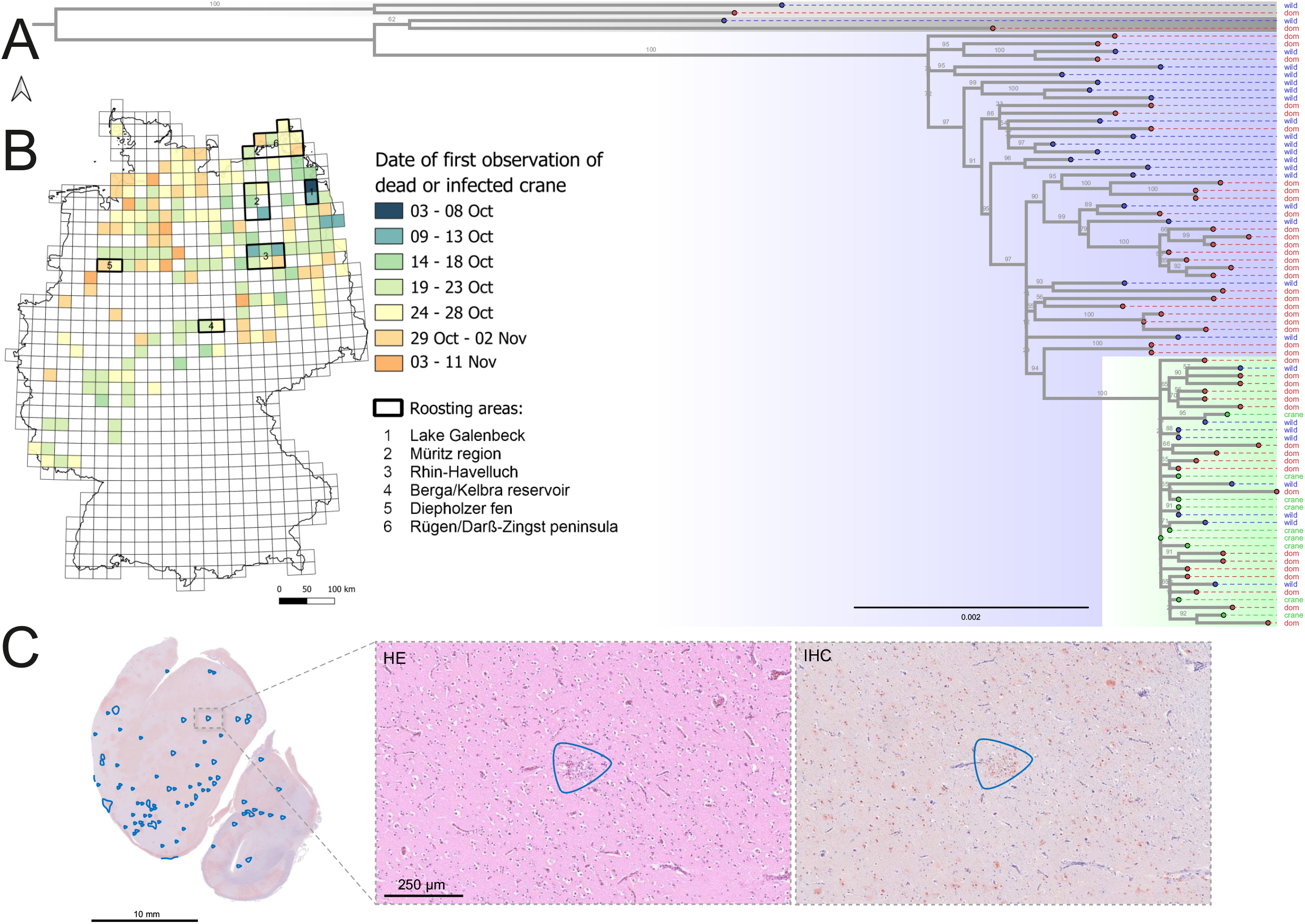
Mass mortality among Eurasian cranes in Germany due to HPAIV H5N1 infections, epidemiology, phylogeny and pathology – (A) Summary of notifications on deceased or infected cranes in Germany, between 3 October and 11 November 2025. Numbers indicate mentioned roosting areas.; (B) Maximum-likelihood tree depicting the HPAIV H5N1 genotype DI.2.1 cluster (blue) comprising poultry (red tip) and wild bird cases (blue tip) from Germany. Sequences from infected cranes (green tip) and their cluster are highlighted (green). For comparison, recent exemplary sequences for previous dominating genotypes DI.2 and DI.1 have been included (grey); (C) Immunohistochemistry (IHC) of brain tissue from HPAIV-infected cranes. Overview of widespread viral antigen (brown staining) with limited inflammatory and necrotic changes (blue shapes). Insets show the corresponding region on consecutive sections, demonstrating abundant antigen-positive cells on IHC but only a small necrotic focus in the matching hematoxylin-eosin-stained section (HE).

The first deceased cranes were found in early October, at Lake Galenbeck, with HPAIV H5N1 confirmed shortly afterwards. To date, no increased mortalities have been reported from regions north (Sweden) and east (Poland and the Baltic states) of the index site. Six cranes associated with the initial outbreak underwent necropsy. Carcasses were well preserved, and in good nutritional and body condition. Gross lesions were consistent with an acute systemic process and dominated by pancreatic necrosis, pulmonary edema, with occasional epicardial and proventricular hemorrhages (Supplement). Three birds were examined histopathologically. Immunohistochemistry revealed the highest influenza A virus-specific antigen loads in the central nervous system (CNS) and pancreas, with moderate labeling in the heart, spleen, and kidneys, and low levels in respiratory and intestinal tissues. Despite widespread viral antigen distribution, necrosis was present only in subsets of antigen-positive areas and did not reflect the full extent of antigen labeling. This pattern, together with the overall limited inflammatory response (Figure 1C; Supplement), supports an acute and rapidly progressive disease process. Reported neurological signs in the field correlate with widespread CNS infection.

HPAIV H5N1 was confirmed by RT-qPCR in oropharyngeal and cloacal swabs and in lung and brain tissues, with very high virus loads (Supplement). Genome sequencing of nine cranes and 25 HPAIV H5N1-positive samples from wild birds, as well as 43 samples from domestic holdings (Supplement), identified HPAIV H5N1 2.3.4.4b of genotype DI.2.1 (Figure 1A). Since September 2025, DI.2.1-like viruses have been detected almost simultaneously at multiple site across Europe [4], appearing to be a sublineage of the Gs/Gd-like HPAIV genotype DI.2 which dominated the European HPAI epizootiology in 2024 and the first half of 2025 [5]. Phylogenetic analyses indicated that all sequences derived from HPAIV H5-positive cranes fall into a monophyletic cluster of DI.2.1 (Figure 1A). The cluster is interspersed with sequences from domestic poultry and other wild bird species from different families, but indicates a direct spread within crane flocks - strengthened by phylogeographic analyses suggesting spread along bird migration routes (Supplement). It is crucial to differentiate this from the majority of German HPAIV cases observed in wild birds and poultry farms in the analyzed time frame, which exhibit a different and more diverse phylogenetic clustering (Figure 1A), pointing to separate events.

Veterinary authorities and bird conservation organizations initiated carcass removal, though efforts varied and were often hindered by the challenging wetland terrain. Removing carcasses has been deemed useful for reducing the incidence of infections in other gregarious species [6]. Despite mitigation attempts, more than 18,000 deceased cranes were reported in Germany during the 2025 autumn. This clearly outnumbered previously reported detections of HPAIV H5, clade 2.3.4.4b, in cranes in Germany in 2020 (H5N8, n=2) and 2023 (H5N1, n=9).

By early November, crane mortality slowed down and finally almost stopped in Germany. As the cranes continued their migration, the virus spread to France and Spain. At Lake du Der-Chantecoq in northeastern France, 4,000– 5,000 dead cranes were recorded until mid-November. Nationwide, an estimated 15,000–20,000 cranes died. In Spain, the estimated number of fatalities is 1,000– 1,500, including 900 around the Gallocanta lagoon. Altogether, the mortality rate along the Western European flyway in autumn 2025 is estimated at approximately 10% of the crane flyway-population (over 420,000 cranes) [7].

## Discussion

The nearly identical, phylogenetically very closely related sequences of HPAIV H5N1 from Eurasian cranes suggest that once DI.2.1 entered the population, cranes became both severely affected hosts and efficient amplifiers, facilitating rapid intra-species-specific spread along the flyway. Close contact at night roosts in shallow waters likely accelerated the transmission of the virus through contaminated surface water, as even minimal environmental viral loads in surface water have been shown to be sufficient for infecting susceptible species [8].

The source species and location of the current genotype DI.2.1 of HPAIV H5N1 remain to be elucidated. Additional sequence data will help identifying HPAIV H5N1 outbreaks in poultry farms as possible sources and/or sinks of the virus and tracing transmission chains. Although cranes dominated in the clinical presentation of the current H5N1 epizootic, the same virus variant was detected in Germany simultaneously in several aquatic water bird species such as grey lag geese, greater white-fronted geese, barnacle geese, European wigeons and mallards (Figure 1A, Supplement). However, compared to previous seasons, mortality in these species was low and sporadic, possibly due to partial population immunity or characteristics of the current H5N1 genotype. Although these species might have been clinically less vulnerable, they were not necessarily protected from virus infection and excretion. In contrast, the crane population appeared to be fully susceptible, likely due to a lack of prior exposure to HPAIV H5 in western flyway populations. Unfortunately, there are no stored serum samples available to confirm this assumption. Furthermore, it remains unclear if HPAIV H5 circulation influenced cranes’ migratory behaviour, e.g. avoiding heavily affected roosting sites.

## Conclusion

As of 17 November 2025, 18,164 dead cranes had been reported in Germany alone. The full demographic impact of this HPAI-associated mass mortality on the western flyway crane population must be studied. This event highlights critical knowledge gaps in our understanding the epidemiology of HPAIV H5 in wild birds. These gaps are primarily due to due to inadequate active monitoring and the lack of serological data needed to assess population immunity. In addition, the networks driving HPAIV H5 transmission and spread within and between wild bird and poultry populations remain still poorly understood. Furthermore, the characteristics of the current DI2.1 genotype require detailed investigation, particularly in wild Anseriformes.

## Acknowledgments

We gratefully acknowledge the support of Diana Parlow and Marco Beerbohm in performing laboratory analyses and pathological examinations. We are indebted to Dr. G. Nowald, Kranichschutz Deutschland gGmbH and all members of the regional working groups of Kranichschutz Deutschland, NABU, for data on migration and mortality of Eurasian cranes. Further numerous volunteers have reported fatalities, collected field data and participated in the removal of carcasses. We express our gratitude to their engagement as well as to the DDA / Federation of German Avifaunists for providing the valuable data on dead and sick cranes from the online portal ornitho.de via its partner Kranichschutz Deutschland. We would like to thank the Institute of Epidemiology for curating the Avian Influenza database in Germany and for granting us access to it.

## Ethical statement

The analyses presented in this study do not require ethical permits in terms of experimental animal trials. The investigations are based on material submitted in the context of animal disease surveillance in Germany.

## References

1. Nadler-Valency, R., Lourie, E., and Sela-Klein, D., Avian Influenza (H5N1) Outbreak Report in Wild Birds – Hula Valley, Israel A Comparison of Two Outbreaks (2021/22 and 2024). Israel Journal of Veterinary Medicine, 2025. 80(2).

2. European Food Safety, A., European Centre for Disease, P., Control, European Union Reference Laboratory for Avian, I., Adlhoch, C., Fusaro, A., et al., Avian influenza overview September-December 2023. EFSA J, 2023. 21(12): p. e8539. DOI: 10.2903/j.efsa.2023.8539.

3. Djurdjevic, B., Petrovic, T., Gajdov, V., Vidanovic, D., Vucicevic, I., Samojlovic, M., and Pajic, M., Natural infection of common cranes (Grus grus) with highly pathogenic avian influenza H5N1 in Serbia. Frontiers in Veterinary Science, 2024. 11. DOI: 10.3389/fvets.2024.1462546.

4. European Food Safety, A., European Union Reference Laboratory for Avian, I., Ducatez, M., Fusaro, A., Gonzales, J.L., Kuiken, T., et al., Unprecedented high level of highly pathogenic avian influenza in wild birds in Europe during the 2025 autumn migration. EFSA J, 2025. 23(11): p. e9811. DOI: 10.2903/j.efsa.2025.9811.

5. European Food Safety, A., European Centre for Disease, P., Control, European Union Reference Laboratory for Avian, I., Alexakis, L., Buczkowski, H., et al., Avian influenza overview December 2024-March 2025. EFSA J, 2025. 23(4): p. e9352. DOI: 10.2903/j.efsa.2025.9352.

6. Knief, U., Bregnballe, T., Alfarwi, I., Ballmann, M.Z., Brenninkmeijer, A., Bzoma, S., et al., Highly pathogenic avian influenza causes mass mortality in Sandwich Tern breeding colonies across north-western Europe. Bird Conservation International, 2024. 34. DOI: 10.1017/S0959270923000400.

7. Nowald, G., ed. Cranes in Germany and Europe: Review of 2024 with special consideration of the weather. Journal der Arbeitsgemeinschaft Kranichschutz Deutschland - Das Kranichjahr 2024/2025, ed. G. Nowald, et al., 2025: NABU-Erlebniszentrum KRANICHWELTEN. 9–11.

