## Supplement for "High pathogenicity avian influenza virus H5N1 (clade 2.3.4.4b) drives mass mortality in Eurasian crane (*Grus grus*) populations in Germany, 2025"

### 1. Results

#### Additional data on migration patterns and mortality in Eurasian crane (*Grus grus*) flocks in Germany

In Germany, large concentrations of Eurasian cranes (*Grus grus*) typically occur at the Pomeranian coast in the first half of October, at the Rhin-Havelluch in the second half of October, and at the Diepholzer fen from the end of October onwards. In 2025, high-pressure weather patterns and northeastern winds caused three large migratory waves (13 and 18/19 October and 6 November), leading to increased crane densities in southern resting areas (Figure 1B).

The temporal and regional distribution of fatalities associated with HPAIV H5N1 mainly followed known migration trajectories. In October 2025, an unusual backward migration toward the northeast was documented, resulting in virus introduction into major stopover sites around Rügen/Darß-Zingst peninsula (Figure 1B, #6), previously unaffected (Supplement). The first carcass in this region was detected on October 19th, 16 days after the index records at Lake Galenbeck (Figure 1B, #1).

The total numbers of dead cranes recorded so far are: Mecklenburg-Western Pomerania: 560; Brandenburg: 4,500 (ca. 2,250 only around the roost site Linum), Saxony-Anhalt ca. 6,000 (5,433 of these at Berga/Kelbra), Thuringia 711, Lower Saxony 6,000 (Figure 1B).

### Pathological findings and histopathological analyses

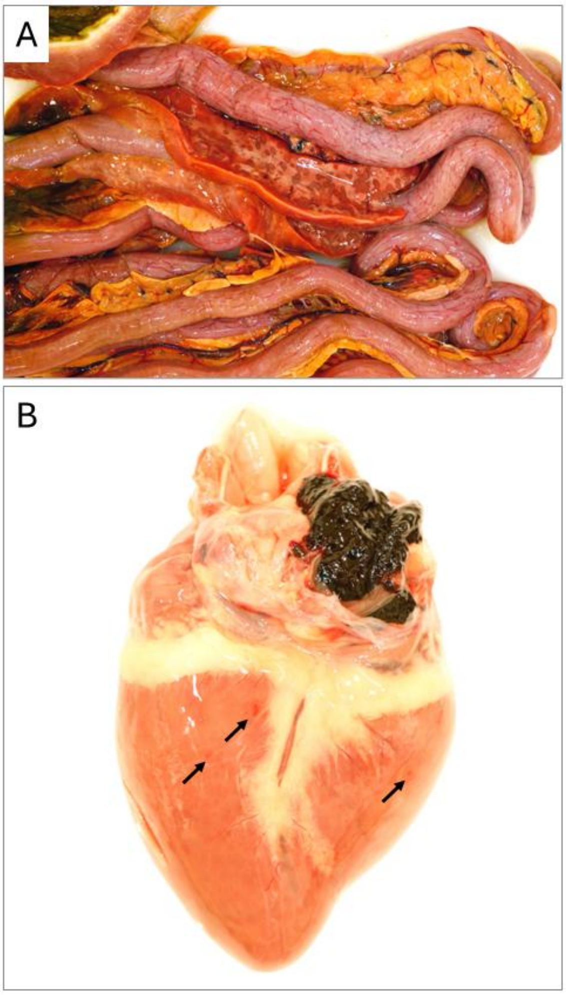


***Supplementary Figure 1*** *Representative gross pathological alterations observed in HPAIV H5N1–infected Eurasian cranes. Marked pancreatic necrosis (A) and epicardial hemorrhage (B) (arrows) are visible.*

***Supplementary Table 1*** *Summary of gross pathological findings in six Eurasian cranes (Grus grus) examined during the initial HPAIV H5N1 outbreak in Germany, autumn 2025. Listed are body condition and the presence (x) or absence (–) of key gross lesions, including pancreatic necrosis, pulmonary edema, epicardial and proventricular hemorrhages, as well as additional findings recorded during necropsy.*

| Bird ID | Body condition | Pancreatic necrosis | Pulmonary edema | Epicardial hemorrhage | Proventricular hemorrhage | Other |
|---|---|---|---|---|---|---|
| #1 | good | x | - | x | x | splenomegaly |
| #3 | good | x | x | - | - | splenomegaly |
| #4 | good | x | x | x | - | splenomegaly, renal congestion |
| #5 | good | x | x | - | - | renal congestion |
| #6 | good | x | x | x | - | renal congestion |
| #7 | good | x | - | - | - | renal congestion |

***Supplementary Table 2*** *Semiquantitative scoring of influenza A viral antigen distribution, necrosis, and inflammation in multiple organs of three Eurasian cranes (Grus grus; #1/#3/#4) naturally infected with HPAIV H5N1. Scores are provided for each organ as antigen score / necrosis score / inflammation score for cranes #1, #3, and #4, respectively. Antigen scores were assigned based on the extent of immunohistochemical (IHC) labelling: 0 = no antigen; 1 = focal/oligofocal (<5%, 1–3 foci; minimal); 2 = multifocal (6–40%, >3 foci; mild); 3 = coalescing (41–80%; moderate); 4 = diffuse (>80%; severe). Necrosis and inflammation were scored on consecutive HE sections using the same scale: 0 = no lesion; 1 = rare (<5%, 1–3 foci; minimal); 2 = multifocal (6–40%, >3 foci; mild); 3 = coalescing (41–80%; moderate); 4 = diffuse (>80%; severe). N/A indicates that the respective tissue was autolytic and could not be reliably assessed.*

*Note: Similar antigen and necrosis scores in some organs reflect the presence of necrotic foci within antigen-positive regions; however, necrosis did not involve all antigen-positive areas.*

| Organ | Antigen score | Necrosis score | Inflammation score |
|---|---|---|---|
| Brain | 3/4/4 | 2/2/2 | 2/2/2 |
| Pancreas | 3/3/3 | 3/3/3 | 2/2/2 |
| Heart | 2/2/2 | 0/0/0 | 0/0/0 |
| Spleen | 2/2/2 | 2/2/2 | 2/1/1 |
| Liver | 1/0/0 | 0/0/0 | 0/0/0 |
| Kidney | 1/1/2 | 1/1/2 | 1/1/2 |
| Lung | 1/1/1 | 0/0/0 | 0/0/0 |
| Proventriculus | 1/1/1 | 0/0/1 | 0/0/0 |
| Gizzard | 0/1/1 | 0/1/1 | 0/1/1 |
| Small intestine | 1/1/1 | N/A | N/A |
| Large intestine | 1/1/1 | N/A | N/A |

### Molecular screenings

***Supplementary Table 3*** *Summary of cycle of threshold (ct) values for oropharyngeal (OPH) and cloacal (CL) swabs, brain and lung tissue from seven Eurasian cranes (Grus grus; #1-#7) that underwent post mortem investigations at the Friedrich-Loeffler-Institut (FLI), Germany. The four blocks represent results for RT-qPCRs targeting different gene fragments of HPAIV H5: matrix gene (M), hemagglutinin subtype H5 (H5), neuraminidase subtype N1 (N1) and the multibasic cleavage site of clade 2.3.4.4b HPAIV (HP). The color coding highlights lower ct values that indicate higher viral loads. For individual #2 no cloacal swab and no lung sample could be taken due to feeding/scavenging damage at the carcass. Asterisks (*) identify those carcasses that had been applied to comprehensive histopathological investigations.*

| M | OPH | CL | Brain | Lung |
|---|---|---|---|---|
| # 1* | 29.06 | 29.93 | 17.1 | 24.09 |
| # 2 | 29.61 | - | 16.3 | - |
| # 3* | 26.9 | 26.66 | 15.93 | 27.16 |
| # 4* | 28.39 | 28.88 | 15.85 | 24.69 |
| # 5 | 30.72 | 31.56 | 22.86 | 26.3 |
| # 6 | 26.18 | 28.43 | 16.81 | 24.28 |
| # 7 | 28.7 | 31.37 | 20.32 | 27.14 |

| N1 | OPH | CL | Brain | Lung |
|---|---|---|---|---|
| # 1* | 25.67 | 26.49 | 14.84 | 20.66 |
| # 2 | 26.18 | - | 12.6 | - |
| # 3* | 23.52 | 23.62 | 12.37 | 23.9 |
| # 4* | 25.02 | 25.43 | 11.12 | 22.02 |
| # 5 | 27.43 | 28.12 | 20.09 | 23.63 |
| # 6 | 23.08 | 25.86 | 12.1 | 21.2 |
| # 7 | 24.93 | 28.39 | 16.59 | 23.71 |

| H5 | OPH | CL | Brain | Lung |
|---|---|---|---|---|
| # 1* | 25.29 | 26.62 | 15.1 | 21.52 |
| # 2 | 25.61 | - | 13.03 | - |
| # 3* | 23.71 | 24.18 | 13.19 | 25.16 |
| # 4* | 25.09 | 26.27 | 12.01 | 22.57 |
| # 5 | 27.82 | 29.16 | 20.72 | 23.95 |
| # 6 | 22.58 | 25.83 | 13.27 | 22.07 |
| # 7 | 25.48 | 29.46 | 17.65 | 24 |

| HP | OPH | CL | Brain | Lung |
|---|---|---|---|---|
| # 1* | 26.05 | 27.45 | 15.25 | 21.58 |
| # 2 | 26.17 | - | 13.29 | - |
| # 3* | 24.05 | 24.3 | 13.26 | 25.3 |
| # 4* | 26 | 26.93 | 12.12 | 22.78 |
| # 5 | 27.99 | 29.31 | 21.08 | 24.24 |
| # 6 | 23.02 | 26.19 | 13.82 | 22.03 |
| # 7 | 26.83 | 29.72 | 17.69 | 24.6 |

### Phylogenetic and phylogeographic analyses

***Supplementary Table 4*** *List of sequenced viruses as applied in Figure 1 A with relevant metadata. Federal states in Germany are listed with their official abbreviations; BB-Brandenburg, BW-Baden-Wuerttemberg, BY-Bavaria, HE-Hesse, MV-Mecklenburg-Western Pomerania, NI-Lower Saxony, NW-North Rhine-Westphalia, SH-Schleswig-Holstein, ST-Saxony-Anhalt, TH-Thuringia.*

| Virus | Federal State | Collection date | host species | host group | genotype |
|---|---|---|---|---|---|
| A/White Stork/Germany-BY/2025AI00446/2025 | BY | 2025-01-14 | *Ciconia ciconia* | wild | DI.1 |
| A/turkey/Germany-NI/2025AI00175/2025 | NI | 2025-01-11 | *Meleagris gallopavo* | domestic | DI.1 |
| A/Eurasian Eagle-Owl/Germany-BY/2025AI03448/2025 | BY | 2025-08-05 | *Bubo bubo* | wild | DI.2 |
| A/chicken/Germany-NI/2025AI02691/2025 | NI | 2025-04-14 | *Gallus gallus domesticus* | domestic | DI.2 |
| A/Eurasian Goshawk/Germany-NW/2025AI06143/2025 | NW | 2025-11-06 | *Accipiter gentilis* | wild | DI.2.1 |
| A/Egyptian Goose/Germany-HE/2025AI05594/2025 | HE | 2025-10-24 | *Alopochen aegyptiaca* | wild | DI.2.1 |
| A/Mallard/Germany-SH/2025AI05309/2025 | SH | 2025-10-24 | *Anas platyrhynchos* | wild | DI.2.1 |
| A/Mallard/Germany-SH/2025AI06081/2025 | SH | 2025-10-28 | *Anas platyrhynchos* | wild | DI.2.1 |
| A/domestic duck/Germany-BB/2025AI04269/2025 | BB | 2025-10-10 | *Anatidae* | domestic | DI.2.1 |
| A/domestic duck/Germany-BB/2025AI04507/2025 | BB | 2025-10-22 | *Anatidae* | domestic | DI.2.1 |
| A/domestic duck/Germany-BB/2025AI04597/2025 | BB | 2025-10-24 | *Anatidae* | domestic | DI.2.1 |
| A/domestic duck/Germany-NI/2025AI04414/2025 | NI | 2025-10-21 | *Anatidae* | domestic | DI.2.1 |
| A/domestic duck/Germany-NI/2025AI05737/2025 | NI | 2025-11-03 | *Anatidae* | domestic | DI.2.1 |
| A/domestic duck/Germany-TH/2025AI04226/2025 | TH | 2025-09-30 | *Anatidae* | domestic | DI.2.1 |
| A/domestic duck/Germany-TH/2025AI04318/2025 | TH | 2025-10-15 | *Anatidae* | domestic | DI.2.1 |
| A/domestic goose/Germany-BB/2025AI04422/2025 | BB | 2025-10-20 | *Anatidae* | domestic | DI.2.1 |
| A/domestic goose/Germany-BY/2025AI04272/2025 | BY | 2025-10-09 | *Anatidae* | domestic | DI.2.1 |

| | | | | | |
|---|---|---|---|---|---|
| A/domestic goose/Germany-NI/2025AI05621/2025 | NI | 2025-11-04 | *Anatidae* | domestic | DI.2.1 |
| A/domestic goose/Germany-NW/2025AI05427/2025 | NW | 2025-11-01 | *Anatidae* | domestic | DI.2.1 |
| A/domestic goose/Germany-NW/2025AI05429/2025 | NW | 2025-11-01 | *Anatidae* | domestic | DI.2.1 |
| A/wild goose/Germany-NI/2025AI04302/2025 | NI | 2025-10-09 | *Anatidae* | wild | DI.2.1 |
| A/wild goose/Germany-SH/2025AI04294/2025 | SH | 2025-10-10 | *Anatidae* | wild | DI.2.1 |
| A/Greylag Goose/Germany-BB/2025AI04273/2025 | BB | 2025-10-09 | *Anser anser* | wild | DI.2.1 |
| A/Greylag Goose/Germany-BY/2025AI05401/2025 | BY | 2025-10-29 | *Anser anser* | wild | DI.2.1 |
| A/Greylag Goose/Germany-MV/2025AI04522/2025 | MV | 2025-10-17 | *Anser anser* | wild | DI.2.1 |
| A/domestic goose/Germany-BB/2025AI04909/2025 | BB | 2025-10-27 | *Anser anser domesticus* | domestic | DI.2.1 |
| A/Bean goose/Germany-MV/2025AI06114/2025 | MV | 2025-11-04 | *Anser fabalis* | wild | DI.2.1 |
| A/Canada Goose/Germany-HE/2025AI06150/2025 | HE | 2025-11-04 | *Branta canadensis* | wild | DI.2.1 |
| A/Barnacle Goose/Germany-NI/2025AI05001/2025 | NI | 2025-10-16 | *Branta leucopsis* | wild | DI.2.1 |
| A/buzzard/Germany-NI/2025AI06173/2025 | NI | 2025-11-04 | *Buteo* | wild | DI.2.1 |
| A/Common Buzzard/Germany-TH/2025AI05077/2025 | TH | 2025-10-25 | *Buteo buteo* | wild | DI.2.1 |
| A/Muscovy Duck/Germany-TH/2025AI04235/2025 | TH | 2025-10-03 | *Cairina moschata* | domestic | DI.2.1 |
| A/Black-headed Gull/Germany-SH/2025AI05322/2025 | SH | 2025-10-25 | *Chroicocephalus ridibundus* | wild | DI.2.1 |
| A/White Stork/Germany-HE/2025AI05330/2025 | HE | 2025-10-27 | *Ciconia ciconia* | wild | DI.2.1 |
| A/swan/Germany-BW/2025AI05762/2025 | BW | 2025-10-31 | *Cygnus* | wild | DI.2.1 |
| A/swan/Germany-SN/2025AI05084/2025 | SN | 2025-10-27 | *Cygnus* | wild | DI.2.1 |
| A/Mute Swan/Germany-BY/2025AI04202/2025 | BY | 2025-09-22 | *Cygnus olor* | wild | DI.2.1 |
| A/chicken/Germany-BB/2025AI04623/2025 | BB | 2025-10-24 | *Gallus gallus domesticus* | domestic | DI.2.1 |
| A/chicken/Germany-BW/2025AI04486/2025 | BW | 2025-10-21 | *Gallus gallus domesticus* | domestic | DI.2.1 |
| A/chicken/Germany-BY/2025AI05958/2025 | BY | 2025-11-05 | *Gallus gallus domesticus* | domestic | DI.2.1 |
| A/chicken/Germany-MV/2025AI04304/2025 | MV | 2025-10-15 | *Gallus gallus domesticus* | domestic | DI.2.1 |
| A/chicken/Germany-MV/2025AI04343/2025 | MV | 2025-10-19 | *Gallus gallus domesticus* | domestic | DI.2.1 |
| A/chicken/Germany-MV/2025AI04345/2025 | MV | 2025-10-19 | *Gallus gallus domesticus* | domestic | DI.2.1 |
| A/chicken/Germany-MV/2025AI05676/2025 | MV | 2025-11-05 | *Gallus gallus domesticus* | domestic | DI.2.1 |
| A/chicken/Germany-NI/2025AI04332/2025 | NI | 2025-10-16 | *Gallus gallus domesticus* | domestic | DI.2.1 |
| A/chicken/Germany-NI/2025AI04500/2025 | NI | 2025-10-22 | *Gallus gallus domesticus* | domestic | DI.2.1 |
| A/chicken/Germany-NI/2025AI05396/2025 | NI | 2025-11-03 | *Gallus gallus domesticus* | domestic | DI.2.1 |
| A/chicken/Germany-NI/2025AI05630/2025 | NI | 2025-11-04 | *Gallus gallus domesticus* | domestic | DI.2.1 |
| A/chicken/Germany-NI/2025AI05634/2025 | NI | 2025-11-04 | *Gallus gallus domesticus* | domestic | DI.2.1 |
| A/chicken/Germany-NW/2025AI05951/2025 | NW | 2025-11-03 | *Gallus gallus domesticus* | domestic | DI.2.1 |
| A/chicken/Germany-SH/2025AI04260/2025 | SH | 2025-10-08 | *Gallus gallus domesticus* | domestic | DI.2.1 |
| A/chicken/Germany-SH/2025AI04293/2025 | SH | 2025-10-14 | *Gallus gallus domesticus* | domestic | DI.2.1 |
| A/chicken/Germany-ST/2025AI04901/2025 | ST | 2025-10-28 | *Gallus gallus domesticus* | domestic | DI.2.1 |
| A/Common Crane/Germany-BB/2025AI04308/2025 | BB | 2025-10-14 | *Grus grus* | crane | DI.2.1 |
| A/Common Crane/Germany-MV/2025AI04305/2025 | MV | 2025-10-10 | *Grus grus* | crane | DI.2.1 |
| A/Common Crane/Germany-MV/2025AI04306/2025 | MV | 2025-10-10 | *Grus grus* | crane | DI.2.1 |
| A/Common Crane/Germany-MV/2025AI04870/2025 | MV | 2025-10-20 | *Grus grus* | crane | DI.2.1 |
| A/Common Crane/Germany-SL/2025AI04530/2025 | SL | 2025-10-22 | *Grus grus* | crane | DI.2.1 |
| A/Common Crane/Germany-ST/2025AI04309/2025 | ST | 2025-10-16 | *Grus grus* | crane | DI.2.1 |
| A/Common Crane/Germany-ST/2025AI04354/2025 | ST | 2025-10-16 | *Grus grus* | crane | DI.2.1 |
| A/Common Crane/Germany-TH/2025AI04342/2025 | TH | 2025-10-17 | *Grus grus* | crane | DI.2.1 |
| A/Common Crane/Germany-TH/2025AI04395/2025 | TH | 2025-10-18 | *Grus grus* | crane | DI.2.1 |
| A/gull/Germany-NI/2025AI05197/2025 | NI | 2025-10-24 | *Laridae* | wild | DI.2.1 |
| A/Eurasian Wigeon/Germany-MV/2025AI04352/2025 | MV | 2025-10-16 | *Mareca penelope* | wild | DI.2.1 |

| | | | | | |
|---|---|---|---|---|---|
| A/Turkey/Germany-BB/2025AI04340/2025 | BB | 2025-10-19 | *Meleagris gallopavo* | domestic | DI.2.1 |
| A/Turkey/Germany-NI/2025AI04289/2025 | NI | 2025-10-13 | *Meleagris gallopavo* | domestic | DI.2.1 |
| A/Turkey/Germany-NI/2025AI04346/2025 | NI | 2025-10-20 | *Meleagris gallopavo* | domestic | DI.2.1 |
| A/Turkey/Germany-NI/2025AI04455/2025 | NI | 2025-10-22 | *Meleagris gallopavo* | domestic | DI.2.1 |
| A/Turkey/Germany-NI/2025AI04604/2025 | NI | 2025-10-24 | *Meleagris gallopavo* | domestic | DI.2.1 |
| A/Turkey/Germany-NI/2025AI04616/2025 | NI | 2025-10-24 | *Meleagris gallopavo* | domestic | DI.2.1 |
| A/Turkey/Germany-NI/2025AI05383/2025 | NI | 2025-11-03 | *Meleagris gallopavo* | domestic | DI.2.1 |
| A/Turkey/Germany-NI/2025AI05388/2025 | NI | 2025-11-03 | *Meleagris gallopavo* | domestic | DI.2.1 |
| A/Turkey/Germany-NI/2025AI05496/2025 | NI | 2025-11-04 | *Meleagris gallopavo* | domestic | DI.2.1 |
| A/Turkey/Germany-NI/2025AI05614/2025 | NI | 2025-11-04 | *Meleagris gallopavo* | domestic | DI.2.1 |
| A/Turkey/Germany-NI/2025AI05625/2025 | NI | 2025-11-04 | *Meleagris gallopavo* | domestic | DI.2.1 |
| A/Turkey/Germany-NW/2025AI04478/2025 | NW | 2025-10-21 | *Meleagris gallopavo* | domestic | DI.2.1 |
| A/Turkey/Germany-SN/2025AI05965/2025 | SN | 2025-11-05 | *Meleagris gallopavo* | domestic | DI.2.1 |
| A/Eurasian Curlew/Germany-RP/2025AI05176/2025 | RP | 2025-10-26 | *Numenius arquata* | wild | DI.2.1 |
| A/Great Cormorant/Germany-TH/2025AI04405/2025 | TH | 2025-10-21 | *Phalacrocorax carbo* | wild | DI.2.1 |
| A/Eurasian Woodcock/Germany-ST/2025AI04948/2025 | ST | 2025-10-22 | *Scolopax rusticola* | wild | DI.2.1 |
| A/Tawny Owl/Germany-BY/2025AI04275/2025 | BY | 2025-10-06 | *Strix aluco* | wild | DI.2.1 |

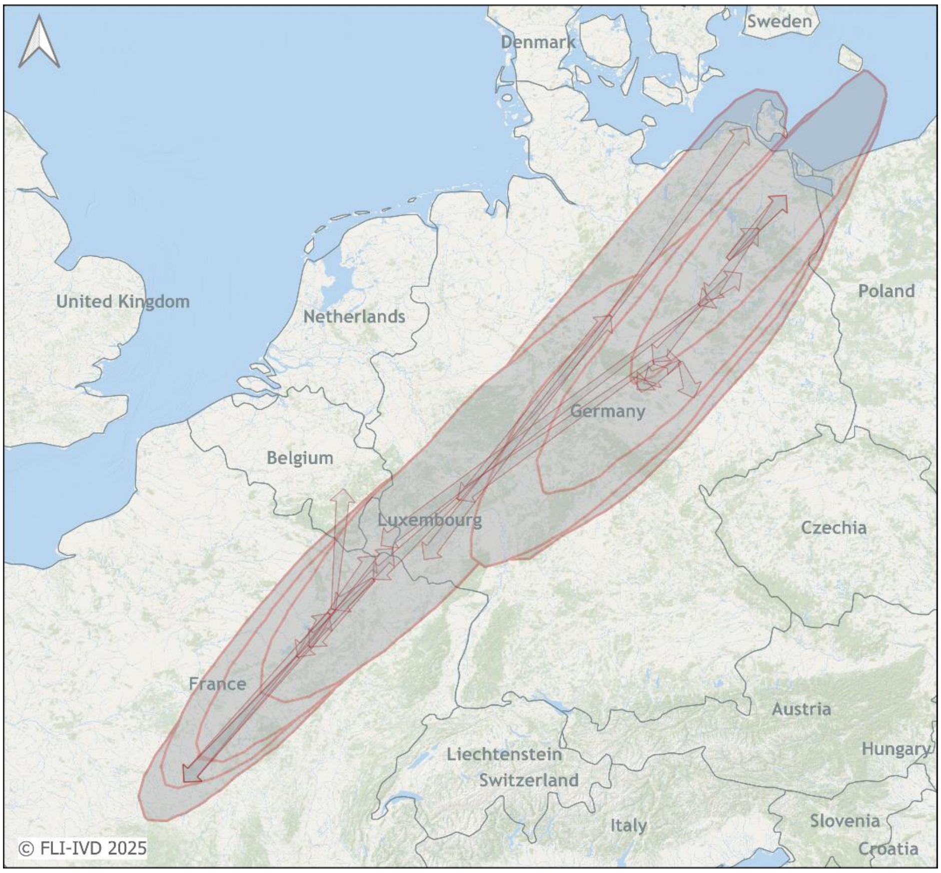


***Supplementary Figure 2*** *Spread of H5N1 HPAI in Eurasian cranes inferred by spatial-time phylogeography with H5 HPAI viral genomes collected from European countries October 2025. Arrows indicated directed spread. Polygons show areas with 95% high posterior densities.*

### 2. Material and Methods

*Summary of report on cranes with clinical signs and mortality in Eurasian cranes*

Reports of deceased or infected cranes between 01.10.2025 and 11.11.2025 (n = 1398) were selected from database at https://www.ornitho.de/ and provided by the Dachverband Deutscher Avifaunisten (DDA). The TK50 grid provided by Bundesamt für Kartographie and Geodäsie (https://www.bkg.bund.de/) was used to demonstrate the timeline of the virus spread in cranes by color-coding each grid cell according to the first report of deceased or infected cranes.

*Necropsy and tissue sampling*

Six common cranes underwent necropsy. CNS, heart, lungs, pancreas, liver, spleen, kidneys, proventriculus, gizzard, small intestine, colon, and caeca were sampled and fixed in 10% neutral-buffered formalin.

*Histopathology*

Formalin-fixed tissues were paraffin-embedded, sectioned at 2–3 µm, and stained with hematoxylin and eosin. Necrosis and inflammation were scored on a 0–4 scale: 0 = no lesion; 1 = minimal (<5%), 2 = mild (6–40%), 3 = moderate (41–80%); 4 = severe (>80%).

*Immunohistochemistry*

IHC was performed using a mouse monoclonal antibody against the influenza A virus matrix (M1) protein (ATCC clone HB-64) [1]. Antigen distribution was scored 0–4: 0 = no antigen; 1 = minimal (<5%, focal-oligofocal); 2 = mild (6–40%, multifocal); 3 = moderate (41–80%, coalescing); 4 = severe (>80%, diffuse).

*Molecular Screenings*

Oropharyngeal and cloacal swabs, as well as tissue samples from brain and lungs, had been screened in RT-qPCRs targeting different genome sections: matrix protein (M1.4), hemagglutinin HA5 (H5), neuraminidase NA1 (N1) and the multibasic cleavage site of clade 2.3.4.4b HPAIV H5 [2, 3].

*Phylogenetic and phylogeographic analyses*

Sequencing of avian influenza-positive samples collected in Germany 2023 was performed using an amplicon-based protocol on nanopore platforms (Oxford Nanopore Technology, Oxford, UK). Briefly, RNA was transcribed into DNA using the Superscript III One-Step and Platinum Taq kit (Thermo Fisher Scientific, USA) with Influenza A specific primers, each binding to the conserved 3' or 5' end of all Influenza RNA segments (Pan-IVA-1F_BsmF: TATTCGTCTCAGGG-AGCRAAAGCAGG; Pan-IVA-1R_BsmR: ATATCGTCTCGTATT-AGTAGAAACAAGG). DNA amplicons were purified with Agencourt AMPure XP magnetic beads (Beckmann Coulter, Krefeld,

Germany) using DNA LoBind Tubes (Eppendorf, Wesseling-Berzdorf, Germany). Quantification of nucleic acids was done with Qubit Fluorometry (Thermo Fisher Scientific, USA). Approximately 200 ng of cDNA was sequenced using a transposase-based library preparation approach with Rapid Barcoding (SQK-RBK114, Oxford Nanopore Technologies, Oxford, UK) and on a PromethION P2 Solo instrument with the latest MinKNOW software core (v6.5.14). High accuracy base calling of the raw data using Dorado (v7.9.8, Oxford Nanopore Technologies) was followed by demultiplexing, a quality check and a trimming step to remove poor quality, adapter and short (<20 bp) sequences. The generated data were stored in FASTQ and POD5 data formats. The bioinformatics software suite Geneious Prime (GraphPad Software LLC, version 2025.1.3) was used for analysis. Sequences were trimmed to remove primer sequences. Consensus sequences were obtained using an iterative map-to-reference approach with Minimap2 (vs 2.24) [4]. Reference genomes were selected from a curated collection of all HA and NA subtypes and a selection of internal gene sequences to cover all potentially circulating viral strains. Polishing of the final genome sequences and annotation was performed manually after consensus generation (threshold matching 60% of bases of total adjusted quality.

Generated sequence data are available in public INSDC databases under accession PX662097 - PX662736. Sequences from other laboratories are listed in DOI 10.55876/gis8.251121ev with the acknowledgement of the originating and submitting laboratories.

Segment-specific and concatenated whole-genome multiple alignments were generated with MAFFT (v7.450) [5] and subsequent maximum likelihood (ML) trees were calculated with RAxML (v8.2.1) [6] using a GTR GAMMA with rapid bootstrapping and search for the best scoring ML tree, supported by 1000 bootstrap replicates, or for large alignments with FastTree (v2.1.11) [7].

Time- scaled trees of concatenated genomes of the same genotype were calculated with the BEAST X (v10.5.0) [8] software package using a GTR GAMMA substitution, an uncorrelated relaxed clock with a lognormal distribution and coalescent constant population tree models. Phylogeographic continuous trait spatial diffusion models were calculated for genotype-based sets using a Bayesian coalescent model with latitude and longitude of the sampling. Chain lengths were set to suitable iterations and convergence checked via Tracer (v1.7.1) [9]. Time- scaled summary maximum clade credibility trees (MCC) with 10% post- burn- in posterior were created using TreeAnnotator (v1.10.4) and visualized with FigTree (V1.4.4). The robustness of the MCC trees was evaluated using 95% highest posterior density confidence intervals at each node and posterior confidence values as branch support. The spatio-temporal diffusion models were analyzed and visualized using Spread (v.1.0.7) [10] and QGIS (v.3.16, QGIS.org). Geographical geojson vector maps were created with open data provided by the Federal Agency for Cartography and Geodesy (http://opendatalab.de/projects/geojson-utilities/) (https://gdz.bkg.bund.de/).

### 3. Reference list for the supplementary material

1. Hohensee, L., Scheibner, D., Schafer, A., Shelton, H., Mettenleiter, T.C., Breithaupt, A., et al., *The role of PB1-F2 in adaptation of high pathogenicity avian influenza virus H7N7 in chickens.* Vet Res, 2024. **55**(1): p. 5. DOI: 10.1186/s13567-023-01257-8.

2. Hassan, K.E., Ahrens, A.K., Ali, A., El-Kady, M.F., Hafez, H.M., Mettenleiter, T.C., et al., *Improved Subtyping of Avian Influenza Viruses Using an RT-qPCR-Based Low Density Array: 'Riems Influenza a Typing Array', Version 2 (RITA-2).* Viruses, 2022. **14**(2): p. 415. DOI: 10.3390/v14020415.

3. Iervolino, M., Günther, A., Begeman, L., Aguado, B., Bestebroer, T.M., Bellido-Martin, B., et al., *The expanding avian influenza panzootic: skua die-off in Antarctica (preprint).* bioRxiv, 2025. DOI: 10.1101/2025.04.25.650384.

4. Li, H., *Minimap2: pairwise alignment for nucleotide sequences.* Bioinformatics, 2018. **34**(18): p. 3094-3100. DOI: 10.1093/bioinformatics/bty191.

5. Katoh, K. and Standley, D.M., *MAFFT multiple sequence alignment software version 7: improvements in performance and usability.* Mol Biol Evol, 2013. **30**(4): p. 772-80. DOI: 10.1093/molbev/mst010.

6. Stamatakis, A., *RAxML version 8: a tool for phylogenetic analysis and post-analysis of large phylogenies.* Bioinformatics, 2014. **30**(9): p. 1312-3. DOI: 10.1093/bioinformatics/btu033.

7. Price, M.N., Dehal, P.S., and Arkin, A.P., *FastTree 2--approximately maximum-likelihood trees for large alignments.* PLoS One, 2010. **5**(3): p. e9490. DOI: 10.1371/journal.pone.0009490.

8. Baele, G., Ji, X., Hassler, G.W., McCrone, J.T., Shao, Y., Zhang, Z., et al., *BEAST X for Bayesian phylogenetic, phylogeographic and phylodynamic inference.* Nat Methods, 2025. **22**(8): p. 1653-1656. DOI: 10.1038/s41592-025-02751-x.

9. Rambaut, A., Drummond, A.J., Xie, D., Baele, G., and Suchard, M.A., *Posterior Summarization in Bayesian Phylogenetics Using Tracer 1.7.* Syst Biol, 2018. **67**(5): p. 901-904. DOI: 10.1093/sysbio/syy032.

10. Bielejec, F., Rambaut, A., Suchard, M.A., and Lemey, P., *SPREAD: spatial phylogenetic reconstruction of evolutionary dynamics.* Bioinformatics, 2011. **27**(20): p. 2910-2. DOI: 10.1093/bioinformatics/btr481.
